# Solution structure and assembly of β-amylase2 from *Arabidopsis thaliana*

**DOI:** 10.1101/751602

**Authors:** Nithesh P. Chandrasekharan, Claire M. Ravenburg, Ian R. Roy, Jonathan D. Monroe, Christopher E. Berndsen

**Affiliations:** Department of Chemistry and Biochemistry, James Madison University, 901 Carrier Dr., MSC 4501, Harrisonburg, VA 22807; Department of Biology, James Madison University, 951 Carrier Dr., MSC 7801, Harrisonburg, VA 22807; Department of Health Sciences, James Madison University, 235 Martin Luther King, Jr Way, MSC 4301, Harrisonburg, VA, United States of America

**Author notes:** To whom correspondence should be addressed., Twitter: @kcat_km.

**Keywords:** amylase, small angle X-ray scattering, Arabidopsis, tetramer

## Abstract

Starch is a key energy storage molecule in plants that requires controlled synthesis and breakdown for effective plant growth. β-amylases (BAMs) hydrolyze starch into maltose to help meet the metabolic needs of the plant. In the model plant, *Arabidopsis thaliana*, there are nine BAMs which have apparently distinct functional and domain structures, although the functions of only a few of the BAMs are known and there are no 3-D structures of BAMs from this organism. Recently, AtBAM2 was proposed to form a tetramer based on chromatography and activity assays of mutants, however there was no direct observation of this tetramer. We collected small-angle X-ray scattering data on AtBAM2 and N-terminal mutants to describe the structure and assembly of the tetramer. Comparison of the scattering of the AtBAM2 tetramer to data collected using the sweet potato (*Ipomoea batatas*) BAM5, which is also reported to form a tetramer, showed there were differences in the overall assembly. Analysis of N-terminal truncations of AtBAM2 identified a loop sequence found only in BAM2 orthologs that appears to be critical for AtBAM2 tetramer assembly as well as activity.

## Introduction

Regulated starch breakdown is important for plant energy metabolism (MacNeill *et al*., 2017). Plants store energy in long, branched glucose chains during the day in a light dependent reaction and then breakdown the starch during the night when light is less available (Zeeman *et al*., 2010; Smirnova *et al*., 2015). Breakdown of starch is, in part, the function of enzymes called β-amylases, which hydrolyze the long dextrin chains of starch into maltose, which is then exported to the cytosol to meet plant metabolic needs (Zeeman *et al*., 2010). Crystal structures of a few β-amylases are known and show that these proteins contain a TIM barrel fold with a pair of catalytic glutamates in the active site required for cleavage of the glycosidic bond (Monroe & Storm, 2018).

β-amylase enzymes (BAMs) are exohydrolases that breakdown starch into maltose by cleaving the α-1,4 glycosidic bonds. However, most of the BAM proteins have not been structurally characterized, including the 9 BAM proteins in the model plant, *Arabidopsis thaliana* (AtBAM1-9) (Monroe & Storm, 2018). Each BAM protein has apparently distinct catalytic activities, structural features, localization, and/or functional regulation (Monroe & Storm, 2018). Recently, AtBAM2 was shown to assemble into a homo-tetramer and required K+ ions for full activity, which are unique properties among characterized BAM proteins (Monroe *et al*., 2017, 2018). While sweet potato (*Ipomoea batatas*) BAM (IbBAM5) was shown to form a tetramer in crystals, in solution this tetramerization is not required for full activity and this enzyme is a homolog of Arabidopsis BAM5 (AtBAM5), which is not known to tetramerize (Monroe & Preiss, 1990; Cheong *et al*., 1995). A BAM protein from *Clostridium thermosulphurogenes* was purified as a tetramer and was active on starch, however biochemical work to support whether this enzyme is required to be in stable tetramer for activity has yet to be done (Shen *et al*., 1988). Thus, the higher order structure of BAM proteins is a relatively unexplored area in this field with implications in cellular function.

Given the proposed importance of the quaternary structure of AtBAM2 for regulation of activity, further exploration of its structure was needed. The initial characterization suggested AtBAM2 formed a tetramer based on SEC-MALS, however direct characterization of the organization and structure was not performed (Monroe *et al*., 2017, 2018). Using small angle X-ray scattering (SAXS), we characterized the solution structure of AtBAM2 and N-terminal truncations of AtBAM2. We confirm that AtBAM2 forms a tetramer in solution and that this tetramer appears distinct from that formed by the IbBAM5. Analysis of the N-terminal truncations revealed a 7-residue sequence at the interface between subunits of the tetramer and appears to be crucial for oligomer formation. Finally, we found that this short sequence is unique to BAM2 orthologs and is conserved across the plant kingdom.

## Methods

### Expression and purification

We expressed *Arabidopsis thaliana* AtBAM2 in E. coli BL21 cells from a pETDuet-1 expression vector with an N-terminal 6-His tag as constructed by (Monroe et al., 2017, 2018). We induced AtBAM2 expression in BL21 E coli at OD_600_ with 0.3 mM IPTG in 2xYT broth with 60 μg/ml ampicillin at 30 °C overnight. Cells were lysed and sonicated in a buffer containing 50 mM NaH_2_PO_4_, pH 8, 500 mM NaCl, and 2 mM imidazole. The supernatant was loaded onto a TALON cobalt column using an AKTA Start and washed with a buffer containing 50 mM HEPES pH 8, 500 mM NaCl, 5% glycerol, 10 mM TCEP, and 40 mM imidazole. The protein was then eluted with buffer containing 50 mM HEPES pH 8, 500 mM NaCl, 5% glycerol, 10 mM TCEP, and 500 mM imidazole. Pure protein, as determined by a band present at ~50 kDa on a 4–20% Tris-Glycine gel stained with Coomassie Blue, was concentrated in a Spin-X UF concentrator with a 5000 MWCO. The concentrated protein was then further purified using a HiLoad 16/60 column filled with Superdex 200 in 50 mM HEPES, pH 7. Pure protein based on SDS-PAGE was concentrated as before and the concentration of protein was determined via absorbance at 280 nm using an extinction coefficient of 94310 M^−1^ cm^−1^ which was calculated from the sequence using ProtParam (Gasteiger et al., 2005). Construction of AtBAM2-Ndel2 was constructed using a similar strategy as that of AtBAM2-Ndel1 (Monroe et al., 2017) but using the primer 5’-TCGAAGAGCGTGATTTTGCGGATCCAGCGTGTGTTCCTGTATATG-3’ and the complementary sequence. AtBAM2 with the Ndel1 or Ndel2 truncations were purified using the same scheme described for the wild-type AtBAM2.

### Homology modeling

Homology modeling of Arabidopsis BAM2 monomer using the plasmid sequence was performed in YASARA Structure (ver 17.6.28) with the default settings. The tetramer was then built by aligning AtBAM2 monomers with the crystallographic tetramer observed in the structure of sweet potato β-amylase (IbBAM5), 1FA2 (Cheong *et al*., 1995). The structure was then equilibrated via molecular dynamics with an AMBER14 force field in explicit solvent at a temperature of 298 K, 0.9% KCl, and pH 7.4 with periodic boundaries (Case *et al*., 2005; Maier *et al*., 2015). Simulations were run with a time step of 100ps and temperature adjusted using a Berendsen thermostat (Krieger *et al*., 2004). Theoretical SAXS data for the homology models were generated using FOXS and plotted using ggplot2 (Wickham, 2010; Schneidman-Duhovny *et al*., 2016). Monomer and dimer models were produced by deleting subunits from the equilibrated model. The Case II tetramer from Cheong, *et al*. was constructed by from the symmetry mates in YASARA followed by energy minimization in YASARA (Cheong *et al*., 1995).

### Small angle X-ray scattering

Data were collected at SiBYLS beamline 12.3.1 at the Advanced Light Source and LBNL. Samples were held at 10 °C during collection. Exposure was 15 seconds with frames collected every 0.3 seconds for 50 total frames per sample. The detector was 2 meters from the sample and the beam energy was 11 keV. Matching buffer exposures were collected before and after samples to ensure there was no difference in the scattering due to contamination of the sample cell. Scattering data were subtracted from buffer and then processed in PRIMUS (ATSAS 2.8.4) to create an average data file (Konarev *et al*., 2003). Data were analyzed using PRIMUS, GNOM, and SCÅTTER (v3.0g) to determine dimensions and create a merged data file from the data sets at 30, 50, and 100 μM AtBAM2 (Konarev *et al*., 2003). We then used the merged data file in DAMMIF (v1.1.2), DAMMIN (v5.3), and GASBOR (v2.3i) to generate the dummy-atom model and aligned the result to the all-atom structure using SASTBX (Svergun, 1992, 1999; Svergun *et al*., 2001; Franke & Svergun, 2009; Liu *et al*., 2012). Dummy atom models from different methods were aligned and visualized using YASARA Structure (Krieger *et al*., 2009). Non-structural plots and statistical analyses were produced in R using the ggplot2 and zoo packages (Wickham, 2010; Zeileis & Grothendieck, 2005).

### Amylase activity assays

Purified wild-type AtBAM2 and the two N-terminal deletions, Ndel1, and Ndel2 AtBAM2 enzymes were tested for amylase activity. Assays were conducted at 25 °C in 0.5 mL containing 50 mM MES (pH 6) and 80 mg/mL soluble starch (Acros Organics). Some assays solution also contained 100 mM KCl. Reactions were stopped after 20 minutes by boiling the reaction tubes in a water bath for 3 minutes. Reducing sugars in each reaction tube were then measured by the Somogyi-Nelson assay with maltose as the standard (Nelson, 1944).

### Sequence alignment

Sequences of BAM proteins from diverse angiosperms and from each major clade of land plants were obtained from NCBI after identification using BLASTp with AtBAM2 as the query. Full-length sequences were aligned using Clustal Omega (Sievers *et al*., 2011) and visualized using BOXSHADE version 3.21.

### Data

SAXS data and models are available online from https://osf.io/eygvm/. The wild-type and Ndel1 scattering data are deposited in the SASBDB as SASDGY4 and SASDGZ4, respectively (Valentini *et al*., 2015).

## Results

### AtBAM2 forms a globular tetramer

AtBAM2 has previously been shown to have an apparently distinct structural organization compared to other characterized BAMs from *Arabidopsis* based on SEC-MALS and mutagenesis (Monroe *et al*., 2018). To provide support for this proposal and to determine the specifics of the structure, we collected small angle X-ray scattering data on AtBAM2. In Figure 1, the data from three concentrations of AtBAM2 are shown along with the calculated radius of gyration (Rg) from each sample shown in the inset. The Rg values are similar for the concentration series and therefore we merged the data for further analysis and modeling.

**Figure 1.**
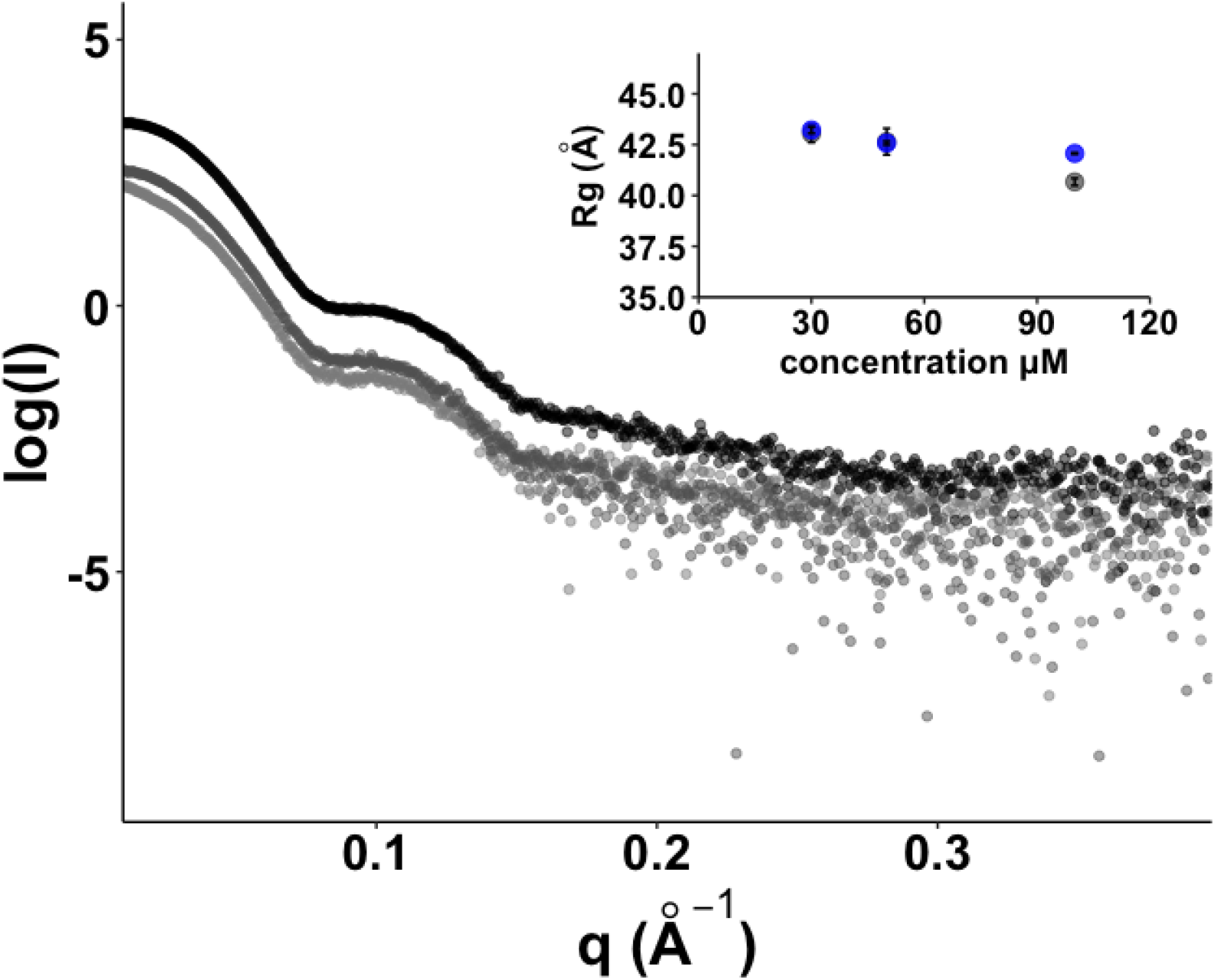
Small angle scattering data for AtBAM2. Inset plot shows the Rg values calculated from the Guinier region (grey dots) or from the P(r) fit (blue dots) using Primus. Error bars show the error from the fit.

Using a merged data set, we calculated the molecular weight of the protein to be 208 kDa [97% Credibility Interval 177 to 221 kDa] (Hajizadeh *et al*., 2018). Further analysis of the structure from the dimensionless Kratky plot suggested a folded, multimeric protein (Rambo & Tainer, 2011). These data are consistent with our previous work using SEC-MALS which suggested that AtBAM2 is tetrameric (Monroe *et al*., 2018). To determine the shape of AtBAM2, we then generated the P(r) plot and then compared the data to several theoretical forms of AtBAM2 along with the existing data on IbBAM5 (Figures 2B, C, and D) (Valentini *et al*., 2015). The elongation ratio of 0.75 calculated from the data suggest a protein that is not spherical, consistent with the Rg value of 42.6 Å and radial cross-section value of 30.4 Å (Figure 2B) (Putnam, 2016). Comparison of the P(r) plot shows that the sweet potato BAM has a distinct profile from AtBAM2 with a broad plateau in r values that could indicate a mixture of tetramers, dimers, or monomers (Figure 2C). Comparison of the P(r) data with theoretical models shows a clear alignment with the tetramer forms of AtBAM2, further supporting the tetrameric structure (Figure 2D). The experimental data are most consistent with a tetramer that is organized as shown in Figure 2E, which is the Case I tetramer from Cheong, *et al*., (Cheong *et al*., 1995). In this structure, the residues 80-93 and 457-485 form a two-fold symmetrical dimer interface. The second interface is formed by amino acids 360-367 and 410-417. These interfaces are compatible with the previous mutagenesis studies on AtBAM2 (Monroe *et al*., 2017, 2018).

**Figure 2.**
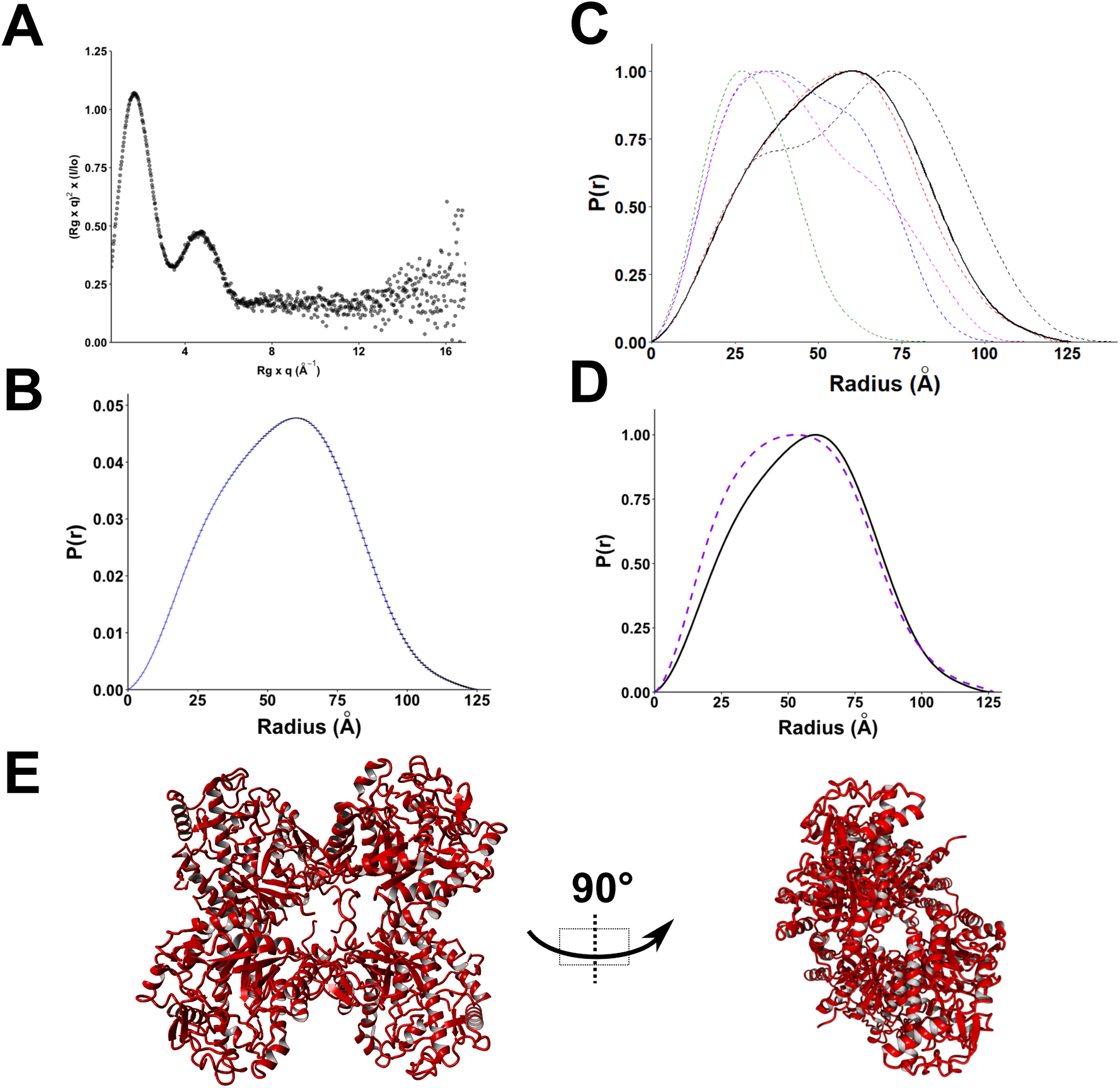
Analysis of the structure of AtBAM2. **(A)** Kratky analysis of merged AtBAM2 scattering data **(B)** P(r) plot of AtBAM2. Dmax was 126 Å, Rg was 42.6 ± 0.03 Å, and the elongation ratio (ER) was 0.75. The total estimate for the fit was 0.79 with an alpha of 39.8. **(C)** Comparison of the AtBAM2 P(r) plot to the theoretical P(r) plots for a tetramer (red or black), dimer (blue or magenta) or a monomer (green). **(D)** Comparison of the AtBAM2 P(r) plot to that of IbBAM5 from SASBDB (SASDA62). **(E)** Model of AtBAM2 tetramer

We next fit the model in Figure 2D to the scattering profile of AtBAM2 using FoXS and the fit matched the shape of the profile with a **χ**^2^ of 8.96 (Figure 3A) (Schneidman-Duhovny *et al*., 2016). We then used model-free analysis to further support and refine our organization of the AtBAM2 tetramer (Franke & Svergun, 2009; Svergun, 1999). Superpositions of the DAMs on the predicted AtBAM2 tetramer from Figure 2D are shown in Figures 3B, C, and D. The resulting dummy-atom models show that AtBAM2 is flattened on two sides but is a largely globular protein in appearance. The TIM barrel portions of AtBAM2 fit inside the dummy-atom model and suggest a hollow or less dense center section. Each dummy atom model has unaccounted for model however there was no consensus across the methods.

**Figure 3.**
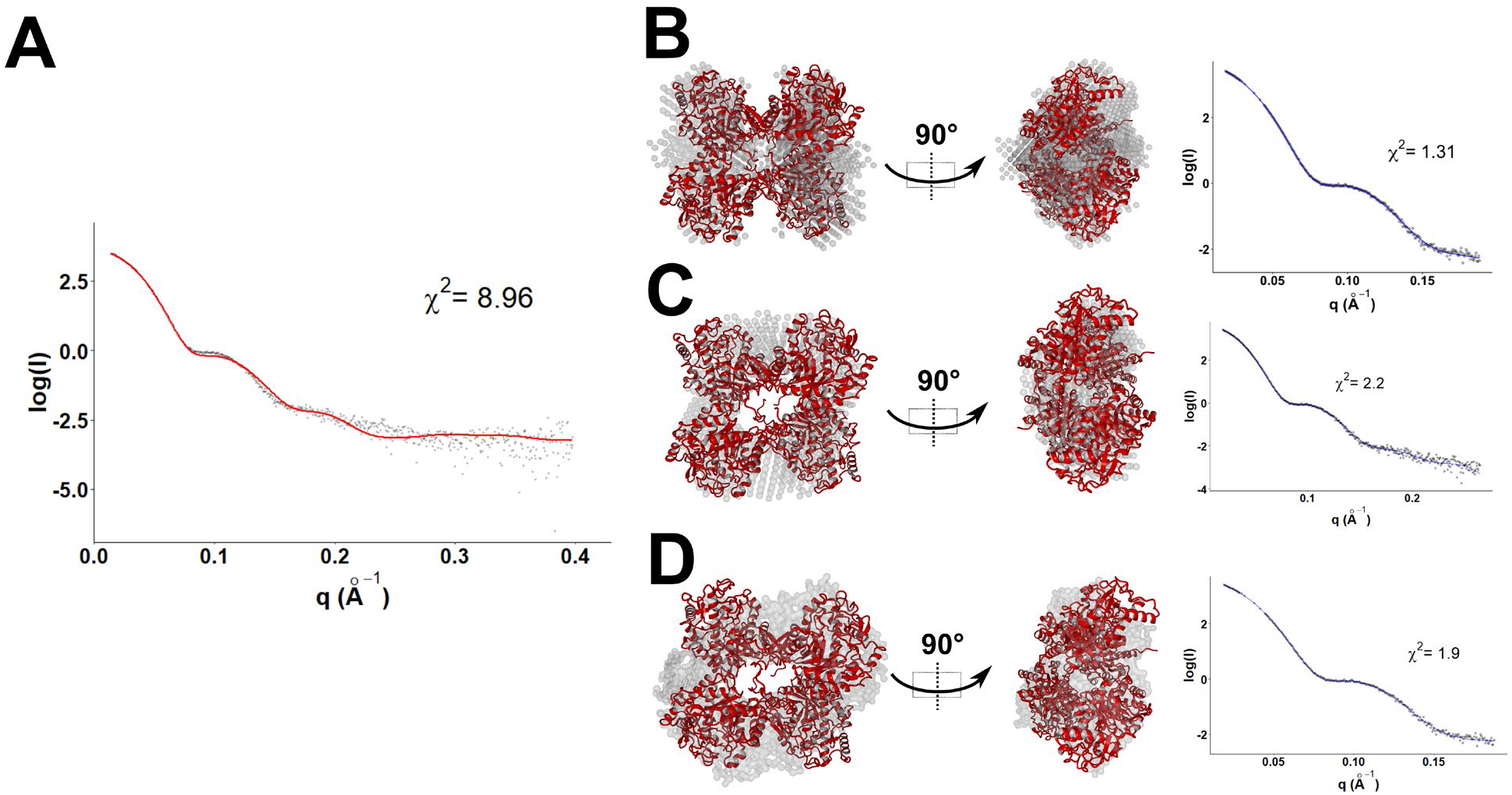
Prediction of AtBAM2 3-D shape. **(A)** Fit of model from Figure 2E to AtBAM2 from the FOXS server. **(B)** DAMMIN, **(C)** DAMMIF, and **(D)** GASBOR dummy atom models based on AtBAM2 data aligned to tetrameric model of BAM2 (red). Fits of the dummy atom models to the data are shown in the right panels of B, C, and D along with χ^2^ of the fit. The NSD of 13 DAMMIF models was 1.299 with a standard deviation of 0.141.

### N-terminal loop is important for AtBAM2 oligomerization

We next analyzed two N-terminal truncations of AtBAM2 to support our structure. Ndel1 removes amino acids 55 to 85 and was biochemically described by Monroe, et al. (2018), while Ndel2 removes 55 to 92 and has not been described before. The difference between these two truncations is 7 residues that are predicted to form a loop on the surface of each subunit and this sequence was found to be conserved among BAM2-like proteins from land plants (Monroe *et al*., 2018). For simplicity we refer to this as the “ERDF loop”. We collected small-angle scattering data on both constructs. Datasets for both proteins showed some degree of aggregation in the sample suggesting some decreased stability (Figure 4A). Kratky analysis and the P(r) distribution of Ndel1 largely align with the shapes observed for the wild-type protein, further indicating that this truncation remains as a tetramer in solution, although we noted some apparently decreased stability (Figures 4A and B).

**Figure 4.**
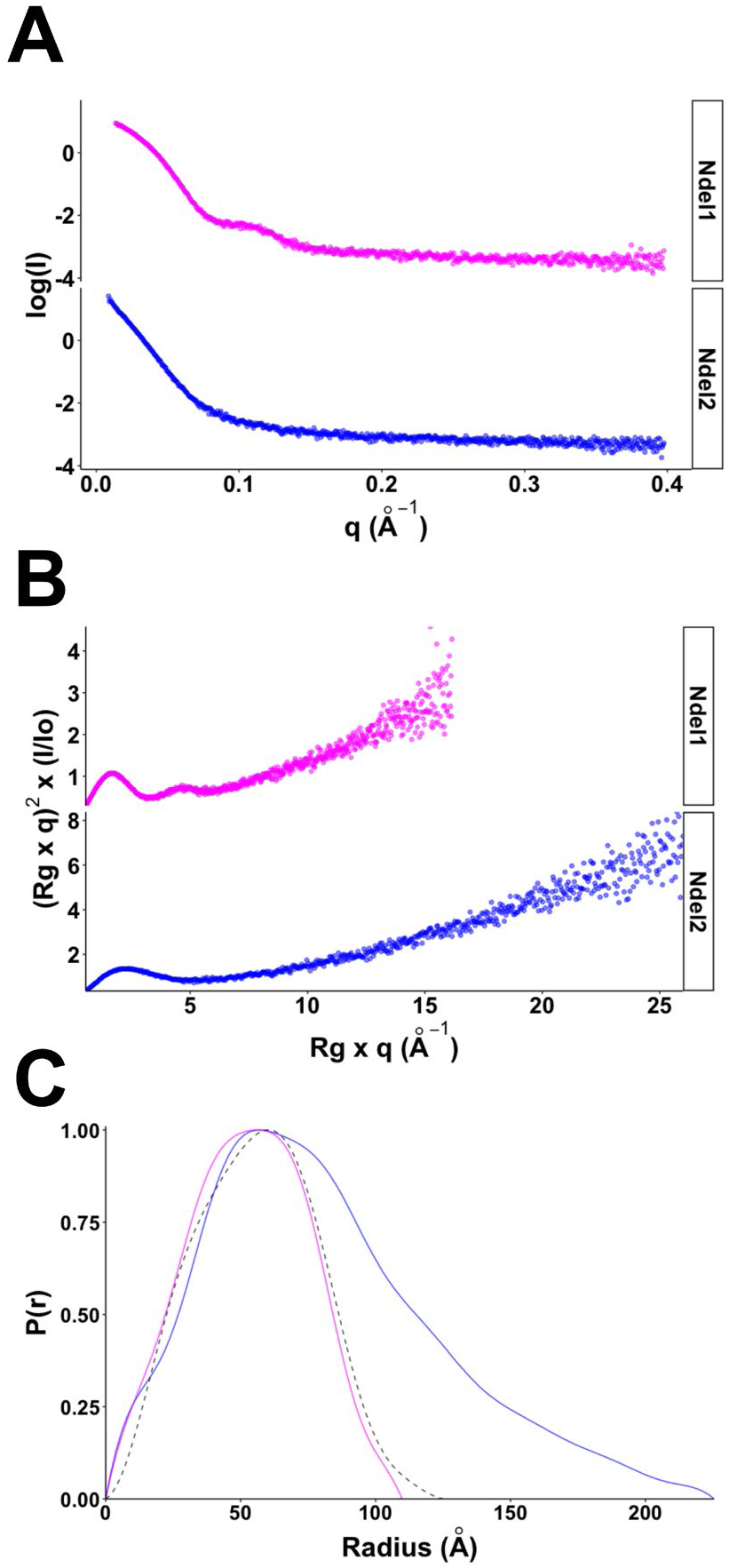
SAXS analysis of AtBAM2 with Ndel1 or Ndel2 truncation. **(A)** Scattering data for Ndel1 (magenta) and Ndel2 (blue). (B) Kratky analysis of Ndel1 and Ndel2. (C) P(r) plot for Ndel1 and Ndel2. The wild-type AtBAM2 data are shown by a dashed black line. For Ndel1, the Dmax was 109.8 Å and the Rg was 40.6 ± 0.05 Å. The total estimate for the fit was 0.84 with an alpha of 2125. For Ndel2, the Dmax was 225.5 Å and the Rg was 65.4 ± 0.4 Å. The total estimate for the fit was 0.75 with an alpha of 6.1.

In contrast to Ndel1, the raw data in Figure 4A suggest that the Ndel2 protein is a mixture of folded and unfolded protein. Like with Ndel1, we observed aggregated protein in the low q region of Figure 4A. The Ndel2 protein was purified and handled alongside the wild-type and Ndel1 proteins and did not show obvious differences during purification, thus we believe the Ndel2 mutation destabilizes the protein. Analysis of data excluding the aggregation showed that the Kratky plot for Ndel2 lacks the second hump seen in the wild-type (Figure 4B). Multiple humps in the Kratky plot is associated with folded multimers, thus these data suggest AtBAM2 with the Ndel2 truncation may be monomeric or unfolded (Rambo & Tainer, 2011; Kikhney & Svergun, 2015). The P(r) plot indicates a larger Dmax than the wild-type with the peak shifted to higher r values which is consistent with an increase in protein disorder (Figure 4C) (Kikhney & Svergun, 2015). All of these data support that residues 86 to 92 of AtBAM2 play a role in forming a stable tetramer.

Molecular modeling of the wild-type and truncated tetramer of AtBAM2 followed by equilibration of the models in an AMBER force field with explicit solvent showed that the truncated tetramer was less stable although the TIM barrel of individual subunits including the active site of AtBAM2 remained largely unchanged (Figure 5A). Sweet potato BAM is known to still be active in the monomeric state, thus we measured the amylase activity of both Ndel proteins and compared these data to the wild-type AtBAM2 (Figure 5B) (Ann *et al*., 1990). Reducing sugar assays show that the NDel2 truncation decreases AtBAM2 activity to ~10% of the wild-type in the presence of KCl and does not show stimulation of activity by KCl. Ndel1 however has activity within two-fold of the wild-type and shows less of a dependence on potassium cations, as observed previously (Monroe *et al*., 2018). These data demonstrate the importance of AtBAM2 oligomerization to the amylase activity of the enzyme.

**Figure 5.**
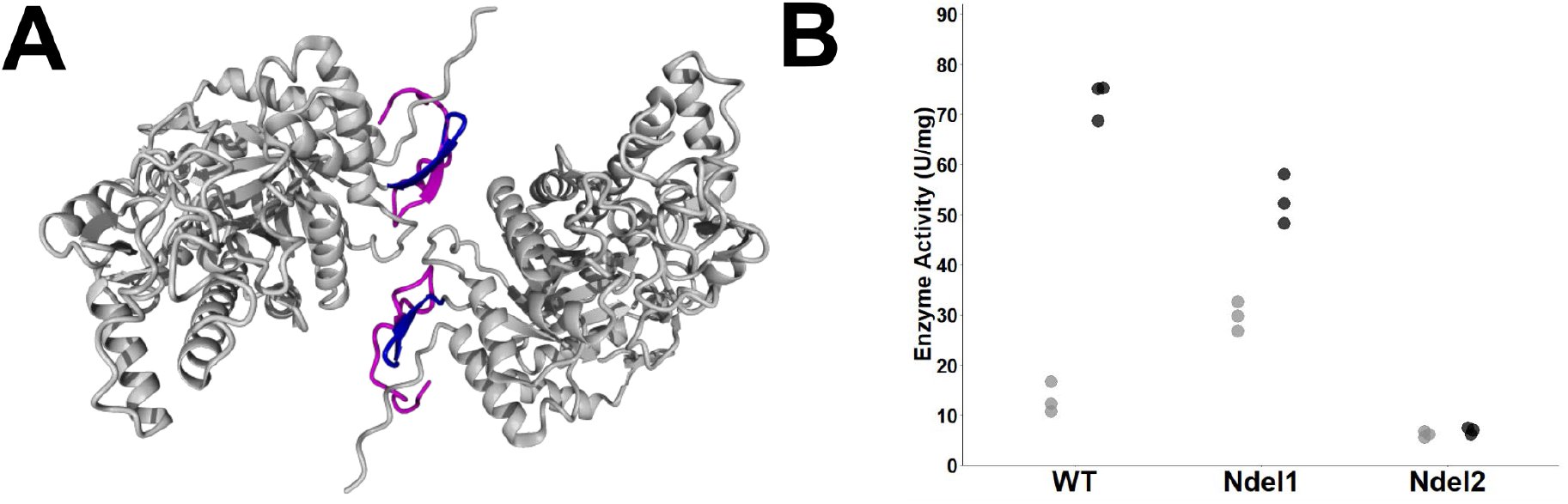
Activity of AtBAM2 with Ndel1 or Ndel2 truncation. **(A)** AtBAM model showing the amino acids removed in the Ndel1 truncation (magenta) and the Ndel2 truncation (magenta and blue). **(B)** Activity of AtBAM2, Ndel1 and Ndel2 truncated enzymes on soluble starch with (black) and without KCl (gray). Each point is an independent measurement with three trials shown for each enzyme.

### Conservation of residues involved in tetramerization

In order to identify BAM2 specific tetramerization residues we aligned sequences of BAM2-like proteins from each major lineage of land plants including nonvascular, seedless vascular, gymnosperm and flowering plants, as well as a BAM2-like protein from *Klebsormidium flaccidum*, a charophyte alga considered to be ancestral to all land plants. Along with these BAM2-like sequences, we also aligned each of the other BAM proteins from Arabidopsis that encode catalytically active proteins and the sweet potato BAM5 sequence (Figure 6). All the BAM proteins align well in the core amylase domain with most of the differences occurring to the N and C-sides of this domain. Residues that were predicted from the tetramer model as occurring at the subunit interfaces and confirmed by mutation as important for catalytic activity were only conserved among all of the BAM2-like proteins and not in BAM1, −3, −5, or −6 from Arabidopsis, or BAM5 from sweet potato (Figure 6). In addition, an active-site adjacent residue, S464, that was found to be essential for cooperativity among the subunits is also conserved among the BAM2-like proteins and not any of the others (Monroe *et al*., 2018). Notably, the ERDF motif is found only in BAM2 orthologs and is not observed in the sweet potato BAM sequence. These alignment data support the structural data and organization of the AtBAM2 tetramer and show that areas apparently critical for oligomer formation are found only in BAM2 orthologs.

**Figure 6.**
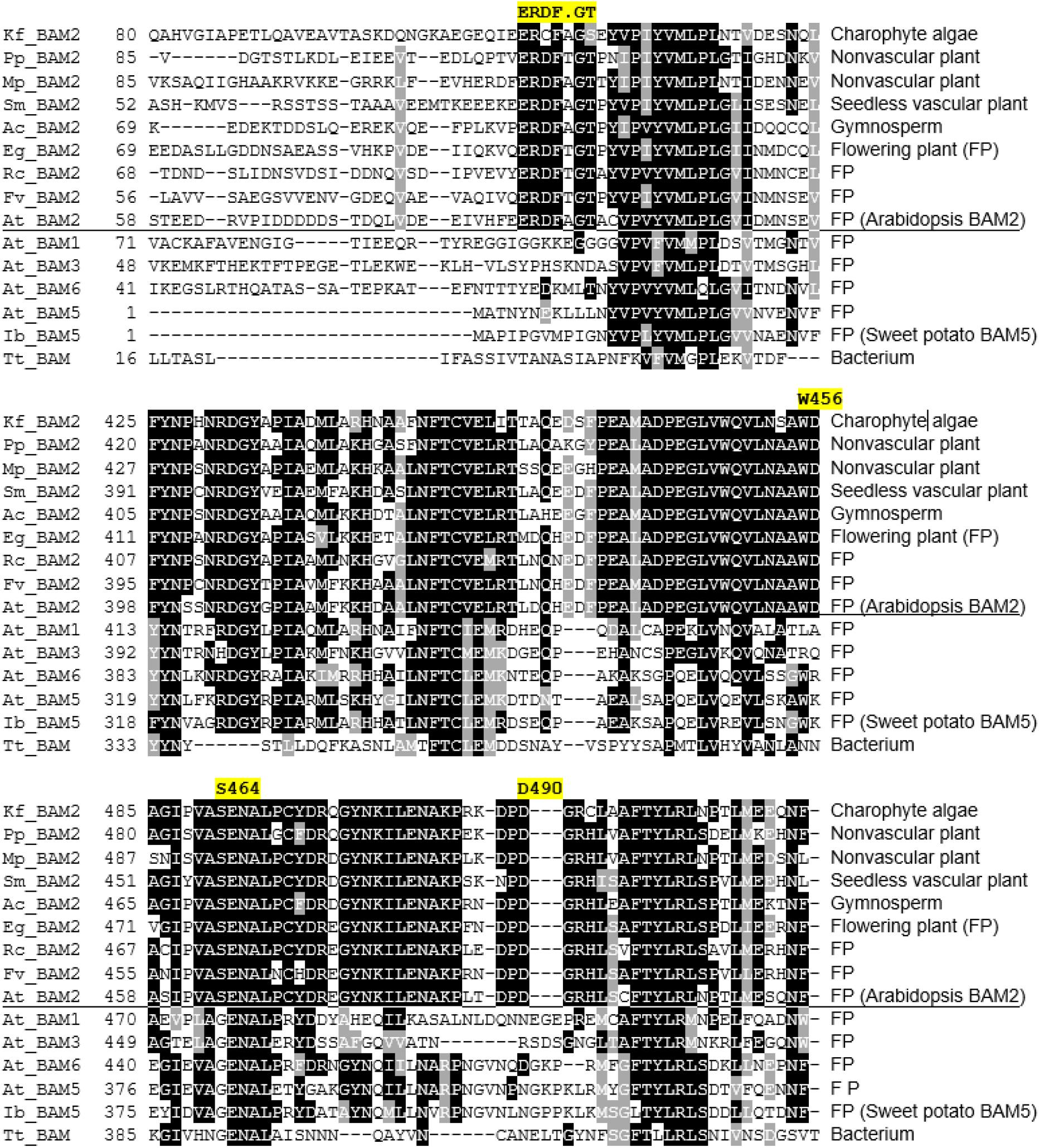
Regions of a sequence alignment of BAM sequences from a charophyte green alga, various land plants, and a bacterium illustrating conservation of residues identified as important for the catalytic activity of Arabidopsis BAM2 (Monroe et al., 2918). Above the horizontal lines are BAM2-like sequences are from the charophyte green alga, Klebsormidium flaccidum (kfl00081_0270), the nonvascular plants Physcomitrella patens (XP_024360526.1) and Marchantia polymorpha (OAE23062.1), the seedless vascular plant Selaginella moellendorffii (XP_024540821.1), the gymnosperm Araucaria cunninghamii (JAG96982.1), and the flowering plants Erythranthe guttata (XP_012843727.1), Ricinus communis (XP_002511858.1), Fragaria vesca (XP_004306786.1) and Arabidopsis thaliana (NP_191958.3). Below the horizontal lines are other catalytically active BAMs from Arabidopsis include BAM1 (NP_189034.1), BAM3 (NP_197368.1), BAM6 (NP_180788.2), and BAM5 (NP_567460.1) as well as BAM5 from sweet potato, Ipomoea batatas (XP_019180769.1) and a BAM from Thermoanaerobacterium thermosulfurigenes (P19584). The full-length sequence alignment is in Figure S1.

## Discussion

*β*-amylases play a crucial role in the degradation of starch during the day-night cycling of plants (Monroe & Storm, 2018). Despite their importance, the structural and functional descriptions of BAM proteins are largely lacking. Recently, we showed that AtBAM2 from Arabidopsis had unique activity requirements and appeared to be tetrameric in SEC-MALS experiments, however the specific organization of the AtBAM2 oligomer was not clear (Monroe *et al*., 2017, 2018). Here, we describe the solution structure of BAM2 from Arabidopsis showing direct evidence for a stable tetramer in solution and how that tetramer is organized. Moreover, we identify a BAM2-specific sequence that is required for stabilizing the tetramer.

While BAM from sweet potato (IbBAM5) has long been reported to form tetramers, the stability of this tetramer is reported to be less than that of AtBAM2 and unlike AtBAM2 is not required for activity (Ann *et al*., 1990; Cheong *et al*., 1995; Costa *et al*., 2016; Monroe *et al*., 2017, 2018). Here, we show that the reference data for IbBAM5 and our data for AtBAM2 do not match exactly and suggest that the IbBAM5 is partially dissociated (Figure 2C). While the tetrameric form for sweet potato BAM5 may be structurally similar to how AtBAM2 organizes into a tetramer, there are clear differences in sequence that lead to IbBAM5 being less stable (Figure 6). One specific difference is the lack of the conserved ERDF loop which is found in the N-terminal extension of all plant BAM2 proteins and is found at one of the interfaces of the AtBAM2 tetramer. This sequence makes contacts with two other regions of the adjacent BAM2 which are conserved in BAM2-like proteins but are not found in other Arabidopsis BAM proteins or in the sweet potato BAM (Figure 6). It has been previously shown that mutations in these regions (W456A, and D490R) that contact the ERDF loop result in decreased activity and changes in the elution volume in SEC-MALS experiments consistent with disruption of the AtBAM2 tetramer (Monroe *et al*., 2018).

The suggestion that the ERDF loop is important for tetramerization was further supported by observing the solution behavior of two mutants which removed the N-terminus up to the ERDF loop (Ndel1) and one that removed the entire N-terminus up to the amylase domain (Ndel2). The Ndel1 protein behaved as the wild-type protein did in solution with scattering data and model-free analysis consistent with a tetrameric protein. The Ndel2 protein however appeared to be partially unfolded in solution and did not appear as a tetramer. Activity assays on both proteins showed that relative to the wild-type, the Ndel2 truncation reduced activity by an order of magnitude and lost the response to potassium ions that the wild-type and Ndel1 proteins show (Figure 5A). Overall, these data support the role of the ERDF loop in stabilizing the tetrameric form of AtBAM2 and the requirement of oligomerization for full AtBAM2 activity.

Sequence alignment of BAM2-like proteins from each major clade of land plants and other BAMs supports the notion that the ERDF loop is important for AtBAM2 tetramerization and function (Figure 6). The N-terminus along with the conserved W456 and D490, which are both essential for function and oligomer assembly, are all located at the interface between two AtBAM2 monomers (Figure 7) (Monroe *et al*., 2018). Moreover, identification of a BAM2-like protein sharing the conserved sequences important for tetramerization in *Klebsormidium flaccidum* suggests that a BAM with the unusual properties of BAM2 predates the origin of land plants and deserves further study to understand its role in starch metabolism.

**Figure 7.**
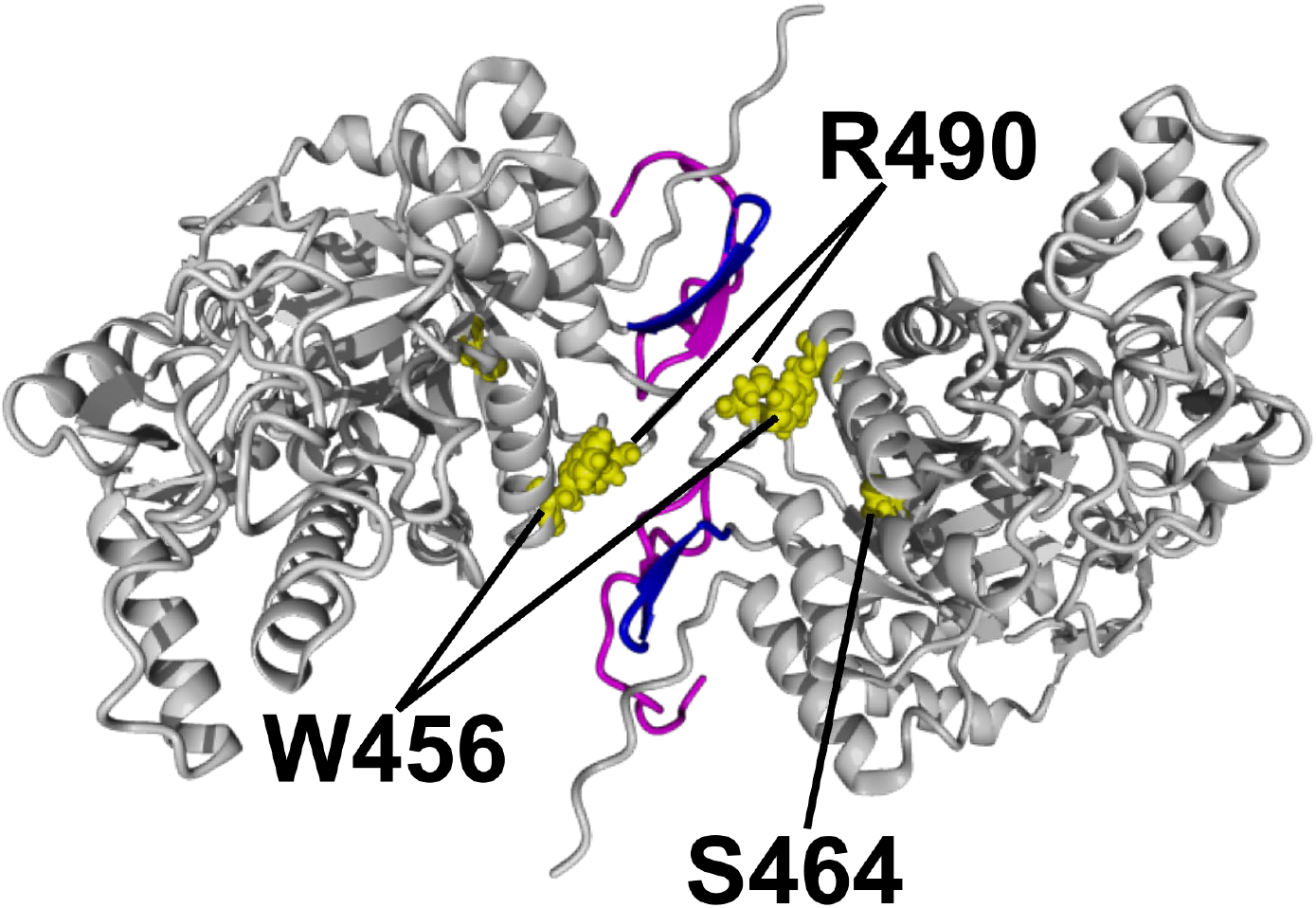
Model of AtBAM2 interface showing the proximity of Ndel1 (magenta) and Ndel2 (magenta and blue) truncation sites to W456, R490, and S464 (shown in yellow).

## Acknowledgements

This work was supported by a National Science Foundation Research at Undergraduate Institutions grant to JM and CEB (MCB-1616467), a Summer Undergraduate Research Fellowship from the American Society of Plant Biologists to CMR, National Science Foundation Research Experience for Undergraduates (CHE-1461175), infrastructure provided by a grant from the Thomas F. and Kate Miller Jeffress Memorial Trust to CEB, and a grant from the 4-VA organization to CEB. We wish to thank Kathryn Burnett and the staff at the SiBYLS beamline for help in collecting scattering data. SAXS data were collected at the Advanced Light Source (ALS), a national user facility operated by Lawrence Berkeley National Laboratory on behalf of the Department of Energy, Office of Basic Energy Sciences, through the Integrated Diffraction Analysis Technologies (IDAT) program, supported by DOE Office of Biological and Environmental Research. Additional support comes from the National Institute of Health project ALS-ENABLE (P30 GM124169) and a High-End Instrumentation Grant S10OD018483. We would also like to thank Jillian Breault for construction of the N-terminal deletion plasmids.

