## Supplemental Figure 1 for "Solution structure and assembly of β-amylase2 from *Arabidopsis thaliana*" (from another version of this preprint: version 1, 2019-08-30)

by

Nithesh P. Chandrasekharan, Claire M. Ravenburg, Ian R. Roy, Jonathan D. Monroe, and Christopher E. Berndsen

**Figure S1.** Full length sequence alignment of BAM sequences from a charophyte green alga, various land plants, and a bacterium illustrating conservation of residues identified as important for the catalytic activity of Arabidopsis BAM2 (Monroe et al., 2918). Above the horizontal lines are BAM2-like sequences are from the charophyte green alga, *Klebsormidium flaccidum* (kfl00081_0270), the nonvascular plants *Marchantia polymorpha* (OAE23062.1) and *Physcomitrella patens* (XP_024360526.1), the seedless vascular plant *Selaginella moellendorffii* (XP_024540821.1), the gymnosperm *Araucaria cunninghamii* (JAG96982.1), and the flowering plants *Erythranthe guttata* (XP_012843727.1), *Ricinus communis* (XP_002511858.1), *Fragaria vesca* (XP_004306786.1) and *Arabidopsis thaliana* (NP_191958.3). Below the horizontal lines are other catalytically active BAMs from Arabidopsis include BAM1 (NP_189034.1), BAM3 (NP_197368.1), BAM6 (NP_180788.2), and BAM5 (NP_567460.1) as well as BAM5 from sweet potato, *Ipomoea batatas* (XP_019180769.1) and a BAM from *Thermoanaerobacterium thermosulfurigenes* (P19584).

```
Kf_BAM2    1 MASRAAVLTS---NASCAALFQSVGALP--DVACKRVAQAVPARSSSQSRFEVQ------
Pp_BAM2    1 MSGL-QLLTVGLVPAKSSEFVGCT---K--FSQCSNLR-LPESRSVQWGVFSHS----WT
Mp_BAM2    1 --------ML---ALTPA---QSG---Q--ERKSRRGG-R-PRSLV-WQHFIVSGERTDK
Sm_BAM2    1 MR-Q-ALLSQ---AAMDARFLCCP---V--ALLPRCHP-------K--------------
Ac_BAM2    1 --------MA---AASSFNLLRCP---N--YNISLRNS-K-TCK-RQFS--SV---KSQG
Eg_BAM2    1 -------MAI---NVSSPELLRRF---T--TLSTRNYA-S-PAFSVAAPLLSVN-SNCRS
Rc_BAM2    1 ---------M---AVSSHQLLHCP---S--PYYS-----T-ASKAQSF-LASIS-TFCCS
Fv_BAM2    1 ---------M---ALLSLQPFRCF---T--PRRP-----S-NVSKTTMPALSFS-PGLWG
At_BAM2    1 ---------M---AIR---LNHSV---I--PVSVK-LG-A-PTRVSARSSLPFS-VGDWR
At_BAM1    1 -------MAL--------NLSHQLGVLAGTPIKSGEMT---DS-----SLLSISPP----
At_BAM3    1 -------MEL--------TLNSSSSLIKRKDAKSSRNQ---ESSSNNMTFAKMKPP----
At_BAM6    1 ------------------------------------------------------------
At_BAM5    1 ------------------------------------------------------------
Ib_BAM5    1 ------------------------------------------------------------
Tt_BAM     1 ------------------------------------------------------------


Kf_BAM2   50 -----TALERSGWKVAPVD----QS------------RISRMSEP---------VQNRGQ
Pp_BAM2   50 HFSPAAISTSGQLREGSKRG-----R----------GVWCRAVEV-------EKLPE---
Mp_BAM2   39 SRRFNVCCKSCGIDMVSRLGTMEKHI----------AFECEKADPD----GRVKIQERLA
Sm_BAM2   30 ------------LAPAARS------N-----------HFVQA-------ITGRKVSN--L
Ac_BAM2   37 SFR----IKSVGLKSSSKK------I-----------HGCQ-SIVLA-LVEEK---G--E
Eg_BAM2   43 IPK------LCKLRESG-------------------------VRSRA-IVGDTELKI--E
Rc_BAM2   36 SSE----FNSCGFRASMVS-------------------FCTRSQNLA-VVRGKK-QR---
Fv_BAM2   37 SPE----FD---------------------------------SRRLR-VVGGGKYRR---
At_BAM2   37 GVS----TFS-G------------------------------ARPLV--LAKVKLRA--E
At_BAM1   34 -----------------SARMMTPKAMNRNYKAHG----TDPSPPMSPILGATRADL--S
At_BAM3   39 -----------------TYQFQ--------------------------------AKN--S
At_BAM6    1 --------------MTSVLGMMNPNLINGRNLHKGSSIFVQDKE------TKKRVQWRLS
At_BAM5    1 ------------------------------------------------------------
Ib_BAM5    1 ------------------------------------------------------------
Tt_BAM     1 --------------------------MIGAFKRLGQKLF-----------------L--T


Kf_BAM2   80 QAHVGIAPETLQAVEAVTASKDQNGKAEGEQIEERCFAGSEYVPIYVMLPLNTVDESNQL
Pp_BAM2   85 -V------DGTSTLKDL-EIEEVT--EDLQPTVERDFTGTPNIPIYVMLPLGTIGHDNKV
Mp_BAM2   85 VKSAQIIGHAAKRVKKE-GRRKLF--EVHERDFERDFAGTTYIPIYVMLPLNTIDENNEV
Sm_BAM2   52 ASH-KMVS---RSSTSS-TAAAVEEMTKEEEKEERDFAGTPYVPIYVMLPLGLISESNEL
Ac_BAM2   69 K------EDEKTDDSLQ-EREKVQE--FPLKVPERDFAGTPYIPVYVMLPLGIIDQQCQL
Eg_BAM2   69 EEDASLLGDDNSAEASS-VHKPVDE--IIQKVQERDFTGTPYVPIYVMLPLGIINMDCQL
Rc_BAM2   68 -TDND--SLIDNSVDSI-DDNQVSD--IPVEVYERDFTGTAYVPVYVMLPLGVINMNCEL
Fv_BAM2   56 -LAVV--SAEGSVVENV-GDEQVAE--VAQIVQERDFTGTPYVPIYVMLPLGVINMNSEV
At_BAM2   58 STEED--RVPIDDDDDS-TDQLVDE--EIVHFEERDFAGTACVPVYVMLPLGVIDMNSEV
At_BAM1   71 VACKAFAVENGIG-----TIEEQR--TYREGGIGGKKEGGGGVPVFVMMPLDSVTMGNTV
At_BAM3   48 VKEMKFTHEKTFTPEGE-TLEKWE--KLH-VLSYPHSKNDASVPVFVMLPLDTVTMSGHL
At_BAM6   41 IKEGSLRTHQATAS-SA-TEPKAT---EFNTTTYEDKMLTNYVPVYVMLQLGVITNDNVL
At_BAM5    1 -----------------------------MATNYNEKLLLNYVPVYVMLPLGVVNVENVF
Ib_BAM5    1 -----------------------------MAPIPGVMPIGNYVPLYVMLPLGVVNAENVF
Tt_BAM    16 LLTASL--------------------IFASSIVTANASIAPNFKVFVMGPLEKVTDF---


Kf_BAM2  140 HNPEQLEKDFAALK-SVGVDGVMCDVWWGLVEREGPGQYDWRGYQELFKRVSGAGLKLQV
Pp_BAM2  135 TNPDDLRQKFNALK-TAEVDGVMVDCWWGLVEGKEPQHYDWSGYRQLFTMVRDCGLKLQV
Mp_BAM2  142 SDPEFLKYDLRALK-SVNVDGVMVDCWWGLVEGKAPQHYNWNGYRQLFNIVREAGLKLQV
Sm_BAM2  107 VNPEGIRRDLQALK-SVNVDGVMVDCWWGLVED-TPQKYNWSAYRELFTIVKESDLKLQV
Ac_BAM2  120 VDPESLQKYLITLK-SANVDGVMVDCWWGIVEGQSPQKYNWSGYRELFDIVCRTKLKLQV
Eg_BAM2  126 VDPDELINHLRVLK-SINVDGVMVDCWWGIVEANAPQQYNWSGYRKLFQIVRDLNLKLQV
Rc_BAM2  122 VDPEGLWNGLKILK-SANVDGVMIDCWWGIVEGNAPQVYDWSGYKRLFQIVNELKLKLQV
Fv_BAM2  110 VEPEVLMDQLRALK-SVGVDGVMVDCWWGIVEAHNPQSYNWSGYKKLFQIVRDLNLKLQV
At_BAM2  113 VEPEELLDQLRTLK-SVNVDGVMVDCWWGIVESHTPQVYNWSGYKKLFQMIRELGLKIQV
At_BAM1  124 NRRKAMKASLQALK-SAGVEGIMIDVWWGLVEKESPGTYNWGGYNELLELAKKLGLKVQA
At_BAM3  104 NKPRAMNASLMALK-GAGVEGVMVDAWWGLVEKDGPMNYNWEGYAELIQMVQKHGLKLQV
At_BAM6   96 ENEESLKKQLKKLK-QSQVDGVMVDVWWGIVESKGPKQYQWSAYRNLFAIVQSFGLKLQA
At_BAM5   32 ADPETLETQLKRLKEEAGVDGVMVDVWWGIIESKGPKQYDWTAYKTLFQLIARLGLKIQA
Ib_BAM5   32 PDKEKVEGELKQVK-AAGCDGVMVDVWWGIIEAKAPKQYDWSAYTELFQLVKKCGLKIQA
Tt_BAM    53 ---NAFKDQLITLK-NNGVYGITTDIWWGYVENAGENQFDWSYYKTYADTVRAAGLKWVP

Kf_BAM2  199 VMSFHQCGGNVGDDTDIPIPPWVHEIAKQNPDIFFTNRKGERNPECLSWGVDKERVLKGR
Pp_BAM2  194 VMSFHQCGGNVGDDVYIPIPQWVLDIGKDNPDIFFTDKSGVVNPECLTWGVDKVRVLRGR
Mp_BAM2  201 VMSFHQCGGNVGDDTNIPIPSWVLDIGEKNPDIFFTDRQGVRNPECLSWGVDKERVLAGR
Sm_BAM2  165 VMSFHQCGGNVGDDVNIPIPKWVLEIGKENPDIFFTDKHGHRNPECLTWGVDKERVLKSR
Ac_BAM2  179 VMSLHECGGNVGDDANITLPKWILQIGEDNPDIFFMDREGQRNKECLTWGIDKERVLRGR
Eg_BAM2  185 VMSFHECGGNVGDDVHIPLPEWVIEIGGKNPDIFFMDRDRRQNRECLTWGIDKERVLRGR
Rc_BAM2  181 VMSFHECGGNVGDDVHIPLPHWVTEIGQTNPDIYFTDREGRRNTECLTWGIGKERVLKGR
Fv_BAM2  169 VMSFHECGGNVGDDVHIPLPYWVTEIGQKNPDIYFTNREGQRIVECLTWGIDKERVLKGR
At_BAM2  172 VMSFHECGGNVGDDVHIQIPEWVREIGQSNPDIYFTDSAGRRNTECLTWGIDKQRVLRGR
At_BAM1  183 VMSFHQCGGNVGDSVTIPLPQWVVEEVDKDPDLAYTDQWGRRNHEYISLGADTLPVLKGR
At_BAM3  163 VMSFHQCGGNVGDSCSIPLPPWVLEEISKNPDLVYTDKSGRRNPEYISLGCDSVPVLRGR
At_BAM6  155 IMSFHRCGGNIGDDVNIPIPKWVLEIGDSNPDIFYTNKSGNRNKECLSLSVDNLSLFRGR
At_BAM5   92 IMSFHQCGGNVGDIVTIPIPQWVRDVGDNDPDIYYTNRKGTRDIEYLSIGVDNLPLFAGR
Ib_BAM5   91 IMSFHQCGGNVGDAVFIPIPQWILQIGDKNPDIFYTNRAGNRNKEYLSLGVDNQRLFQGR
Tt_BAM   109 IMSTHACGGNVGDTVNIPIPSWVWTKDTQ-DNMQYKDEAGNWDNEAVSP------WYSG-


Kf_BAM2  259 TGLEVYYDFMRSFRSAMDEYFSKGVITEVEVGLGACGELRYPSYPERHG-WKYPGIGEFQ
Pp_BAM2  254 TALEVYYDYMRSFRQEMDEFFMDKTITEIEIGLGACGELRYPSYPETRG-WKYPGIGEFQ
Mp_BAM2  261 TGLEIYYDFMRSFRSQFDEFFEDGTITEIEVGLGACGELRYPSYPERHG-WKYPGIGEFQ
Sm_BAM2  225 TGLEVYFDYMRSFRQEFDDLFQSGVIAEIEVGLGACGELRYPSYVPRSG-WEYPGIGEFQ
Ac_BAM2  239 TGLEVCFDYMRSFRMEFNDYFEDGIISEIEIGLGPCGELRYPSYPERHG-WKYPGIGQFQ
Eg_BAM2  245 TALEVYFDYMRSFRVEFDEFFEDGVITEIEIGLGPCGELRYPSYPAKHG-WKYPGIGEFQ
Rc_BAM2  241 TAVEVYFDYMRSFRVEFDEFFEDGMISEIEVGLGPCGELRYPSYPAKHG-WRYPGIGEFQ
Fv_BAM2  229 TAVEVYFDYMRSFRVEFDEFFAEGIISEIEVGLGPCGELRYPSYPAQHG-WKYPGIGEFQ
At_BAM2  232 TALEVYFDYMRSFRVEFDEFFEEKIIPEIEVGLGPCGELRYPSYPAQFG-WKYPGIGEFQ
At_BAM1  243 TPVQCYADFMRAFRDNFKHLL-GETIVEIQVGMGPAGELRYPSYPEQEGTWKFPGIGAFQ
At_BAM3  223 TPIQVYSDFMRSFRERFEGYI-GGVIAEIQVGMGPCGELRYPSYPESNGTWRFPGIGEFQ
At_BAM6  215 TAVEMYRDYMKSFRENMEDFISSGVIIDIEVGLGPAGELRYPSYSETQG-WVFPGIGEFQ
At_BAM5  152 TAVQLYSDYMSSFKENMADLIEAGVIVDIEVGLGPAGELRYPSYPQSQG-WVFPGIGEFQ
Ib_BAM5  151 TALEMYRDFMESFRDNMAEFLKGGDIVDIEVGCGAAGELRYPSYPETQG-WVYPGIGEFQ
Tt_BAM   161 -LTQLYNEFYSSFASNFSSY--KDIITKIYISGGPSGELRYPSYNPSHG-WTYPGRGSLQ


Kf_BAM2  318 CYDKYMMAKLTKAAEA------RNHADWA-HGPEDAGEYNSQPQDTGFFR-DGGNFDSYY
Pp_BAM2  313 CYDKYLLEDLRKAAEA------RGHSHWT-KPPSNAGEYNSRPQDTEFFR-DGGDYDSYY
Mp_BAM2  320 VYDKYLQKSLTKAAVQ------RGHGDWA-RAPDNAGEYNSRPQDTGFFR-DGGDYDSYY
Sm_BAM2  284 CYDKYLIKSLKQAAEA------RGHTEWG-CCPDNAGEYNSKPQETDFFR-DGGDYDSYY
Ac_BAM2  298 CYDKYLQKNLRKAAEA------RGHPFWG-RGPDNAGHYNSRPQDTGFFC-DGGDYDSYY
Eg_BAM2  304 CYDKYLMKSLKKASEV------RGHSFWG-EGPENAGTYNCKPQETKFFS-DGGDYDSYY
Rc_BAM2  300 CYDKYLMRSLSKAAEA------RGHSFWA-RGPDNAGFYNSAPHETGFFR-DGGDYDSYY
Fv_BAM2  288 CYDRYLMENLRKAAEV------RGHSFWA-RAPDNAGSYNFRPQDTGFFR-DGGDYDSYY
At_BAM2  291 CYDKYLMNSLKEAAEV------RGHSFWG-RGPDNTETYNSTPHGTGFFR-DGGDYDSYY
At_BAM1  302 CYDKYSLSSLKAAAET------YGKPEWGSTGPTDAGHYNNWPEDTQFFKKEGGGWNSEY
At_BAM3  282 CYDKYMKSSLQAYAES------IGKTNWGTSGPHDAGEYKNLPEDTEFFRRD-GTWNSEY
At_BAM6  274 CYDKYLRSDYEEEVRR------IGHPEWK--LPENAGEYNSVPGETEFFEYSNGTYLKEE
At_BAM5  211 CYDKYLKKDFKEAAAK------AGHPEWD--LPEDAGEYNDKPEETGFFK-KDGTYVSEK
Ib_BAM5  210 CYDKYMVADWKEAVKQ------AGHADWE--MPGNAGTYNDTPDKTGFFK-PNGTYKTDF
Tt_BAM   217 CYSKAAITSFQNAMKSKYGTIAAVNSAWGTSLT-DFSQISPPTDGDNFFT---NGYKTTY


Kf_BAM2  370 GRWFLKWYSQQLIEHGDRVLHLADMVFEG-T--KIAAKVSGIHWWYKT--ASHAAELCAG
Pp_BAM2  365 GRFFLKWYSDVLIQHGDRVLTFANIAFEG-V--KIAAKVSGIHWWYKT--ASHAAELAAG
Mp_BAM2  372 GRFFLKWYSQVLIEHGDNVLTLANLAFEG-Y--KIAAKVSGIHWWYKT--ASHAAELAAG
Sm_BAM2  336 GRFFLKWYSKVLIEHGDRVLMLAKLAFEG-F--HIASKVSGIHWWYKT--ASHAAELAAG
Ac_BAM2  350 GRFFLRWYSQVLLDHGDRVLSLANLAFEG-T--RIAAKVSGIHWWYKT--ASHAAELAAG
Eg_BAM2  356 GRFFLNWYSQVLIDHGDRVLTFANLAFEG-T--PIAAKLSGIHWWYKT--ASHAAELTAG
Rc_BAM2  352 GRFFLNWYSRVLIDHGDRVLALANLAFEG-T--CISAKVSGIHWWYKT--ASHAAELTAG
Fv_BAM2  340 GRFFLNWYSNVLVDHGDRVLTLANLAFEG-T--SIAAKLSGIHWWYKT--ASHAAELTAG
At_BAM2  343 GRFFLNWYSRVLIDHGDRVLAMANLAFEG-T--CIAAKLSGIHWWYKT--ASHAAELTAG
At_BAM1  356 GDFFLSWYSQMLLDHGERILSSAKSIFEN-MGVKISVKIAGIHWHYGT--RSHAPELTAG
At_BAM3  335 GKFFMEWYSGKLLEHGDQLLSSAKGIFQG-SGAKLSGKVAGIHWHYNT--RSHAAELTAG
At_BAM6  326 GNFFLSWYSKKLLLHGDQILDEANKVFLG-CKLKIAAKVSGIHWWYKT--ESHAAELTAG
At_BAM5  262 GKFFMTWYSNKLIFHGDQILGEANKIFAG-LKVNLAAKVSGIHWLYNH--HSHAAELTAG
Ib_BAM5  261 GKFFLTWYSNNLIIHGDQVLEQANKVFVG-FRVNIAAKVSGIHWWYNH--VSHAAELTAG
Tt_BAM   273 GNDFLTWYQSVLTNELANIASVAHSCFDPVFNVPIGAKIAGVHWLYNSPTMPHAAEYCAG

Kf_BAM2  425 FYNPHNRDGYAPIADMLARHNAAFNFTCVELITTAQEDSFPEAMADPEGLVWQVLNSAWD
Pp_BAM2  420 FYNPANRDGYAAIAQMLAKHGASFNFTCVELRTLAQAKGYPEALADPEGLVWQVLNAAWD
Mp_BAM2  427 FYNPSNRDGYAPIAEMLAKHKAALNFTCVELRTSSQEEGHPEAMADPEGLVWQVLNAAWD
Sm_BAM2  391 FYNPCNRDGYVEIAEMFAKHDASLNFTCVELRTLAQEEDFPEALADPEGLVWQVLNAAWD
Ac_BAM2  405 FYNPSNRDGYAAIAQMLKKHDTALNFTCVELRTLAHEEGFPEAMADPEGLVWQVLNAAWD
Eg_BAM2  411 FYNPANRDGYAPIASVLKKHETALNFTCVELRTMDQHEDFPEALADPEGLVWQVLNAAWD
Rc_BAM2  407 FYNPSNRDGYAPIAAMLNKHGVGLNFTCVEMRTLNQNEDFPEALADPEGLVWQVLNAAWD
Fv_BAM2  395 FYNPCNRDGYTPIAVMFKKHAAALNFTCVELRTLNQHEDFPEAMADPEGLVWQVLNAAWD
At_BAM2  398 FYNSSNRDGYGPIAAMFKKHDAALNFTCVELRTLDQHEDFPEALADPEGLVWQVLNAAWD
At_BAM1  413 YYNTRFRDGYLPIAQMLARHNAIFNFTCIEMRDHEQP---QDALCAPEKLVNQVALATLA
At_BAM3  392 YYNTRNHDGYLPIAKMFNKHGVVLNFTCMEMKDGEQP---EHANCSPEGLVKQVQNATRQ
At_BAM6  383 YYNLKNRDGYRAIAKIMRRHHAILNFTCLEMKNTEQP---AKAKSGPQELVQQVLSSGWR
At_BAM5  319 YYNLFKRDGYRPIARMLSKHYGILNFTCLEMKDTDNT---AEALSAPQELVQEVLSKAWK
Ib_BAM5  318 FYNVAGRDGYRPIARMLARHHATLNFTCLEMRDSEQP---AEAKSAPQELVREVLSNGWK
Tt_BAM   333 YYNY------STLLDQFKASNLAMTFTCLEMDDSNAY--VSPYYSAPMTLVHYVANLANN


Kf_BAM2  485 AGIPVASENALPCYDRQGYNKILENAKPRK-DPD---GRCLAAFTYLRLNPTLMEEQNF-
Pp_BAM2  480 AGISVASENALGCFDRQGYNKILENAKPEK-DPD---GRHLVAFTYLRLSDELMKEHNF-
Mp_BAM2  487 SNISVASENALPCYDRDGYNKILENAKPLK-DPD---GRHLVAFTYLRLNPTLMEDSNL-
Sm_BAM2  451 AGIYVASENALPCYDRDGYNKILENAKPSK-NPD---GRHISAFTYLRLSPVLMEEHNL-
Ac_BAM2  465 AGIPVASENALPCFDRDGYNKILENAKPRN-DPD---GRHLEAFTYLRLSPTLMEKTNF-
Eg_BAM2  471 VGIPVASENALPCYDREGYNKILENAKPFN-DPD---GRHLSAFTYLRLSPDLIEERNF-
Rc_BAM2  467 ACIPVASENALPCYDREGYNKILENAKPLE-DPD---GRHLSVFTYLRLSAVLMERHNF-
Fv_BAM2  455 ANIPVASENALNCHDREGYNKILENAKPRN-DPD---GRHLSAFTYLRLSPVLLERHNF-
At_BAM2  458 ASIPVASENALPCYDREGYNKILENAKPLT-DPD---GRHLSCFTYLRLNPTLMESQNF-
At_BAM1  470 AEVPLAGENALPRYDDYAHEQILKASALNLDQNNEGEPREMCAFTYLRMNPELFQADNW-
At_BAM3  449 AGTELAGENALERYDSSAFGQVVATN-------RSDSGNGLTAFTYLRMNKRLFEGQNW-
At_BAM6  440 EGIEVAGENALPRFDRNGYNQIILNARPNGVNQDGKP--RMFGFTYLRLSDKLLNEPNF-
At_BAM5  376 EGIEVAGENALETYGAKGYNQILLNARPNGVNPNGKPKLRMYGFTYLRLSDTVFQENNF-
Ib_BAM5  375 EYIDVAGENALPRYDATAYNQMLLNVRPNGVNLNGPPKLKMSGLTYLRLSDDLLQTDNF-
Tt_BAM   385 KGIVHNGENALAISNNN---QAYVN------CANELTGYNFSGFTLLRLSNIVNSDGSVT


Kf_BAM2  540 KEFARFVKRLHGEAVDDLNT----------------------------------------
Pp_BAM2  535 KEFSRFVKRLHGKPVESD------------------------------------------
Mp_BAM2  542 DEFALFVKRLHGKQTHFSS-----------------------------------------
Sm_BAM2  506 HEFARFVKRLHGEGV---------------------------------------------
Ac_BAM2  520 YEFERFVKRMHGEAVLDLQA----------------------------------------
Eg_BAM2  526 VEFEQFVKRMHGDSVPDLEHC---LEEEAR-----TVDEKT-------------------
Rc_BAM2  522 IEFERFVKRMHGDLSLSPLLL---LGFSLY-----FLIQLTI-RM----------VCWHA
Fv_BAM2  510 MEFERFVKKMHGEAATDLQIN---PNEENT-----ADSSANL-Q----------------
At_BAM2  513 KEFERFLKRMHGEAVPDLGLA---PGTQET-----NPE----------------------
At_BAM1  529 GKFVAFVKKMGEGRDSHRCREEVEREAEHFVHVTQPLVQ-----EAAVALTH--------
At_BAM3  501 QQLVEFVKNMKEGGHGRRLSKEDTTGSDLYVGFVKGKIAENV-EEAALV-----------
At_BAM6  497 STFKMFLKRMHANQEYCSEPE---RYNHELLPLERSRNDESL-EMFMEETEPFDPFPWLD
At_BAM5  435 ELFKKLVRKMHADQDYCGDAA---KYGHEIVPLKTSNSQLTL-EDIADAAQPSGAFKWDS
Ib_BAM5  434 ELFKKFVKKMHADLDPSPNAI-------SPAVLERSNSAISI-DELMEATKGSRPFPWYD
Tt_BAM   436 SEMAPFVINIVTLT----------PNGTIPVTFTINNATTYYGQNVYIVGSTSDLGNWNT


Kf_BAM2      ------------------------------------------------------------
Pp_BAM2      ------------------------------------------------------------
Mp_BAM2      ------------------------------------------------------------
Sm_BAM2      ------------------------------------------------------------
Ac_BAM2      ------------------------------------------------------------
Eg_BAM2      ------------------------------------------------------------
Rc_BAM2  563 MAYDIKICRCLEIPVYCSYLVCQLRFHFTLW--VHLHDRCGCVWVVSGS-----------
Fv_BAM2      ------------------------------------------------------------
At_BAM2      ------------------------------------------------------------
At_BAM1      ------------------------------------------------------------
At_BAM3      ------------------------------------------------------------
At_BAM6  553 ET-DMSIR------PFESVL-SLLRSTFL-------RKKS--------------------
At_BAM5  491 ET-DLKVD------G---------------------------------------------
Ib_BAM5  486 VT-DMPVD------GSNPFD----------------------------------------
Tt_BAM   486 -TYARGPASCPNYPTWTITL-NLLPGEQIQFKAVKIDSSGNVTWEGGSNHTYTVPTSGTG

Kf_BAM2      --------
Pp_BAM2      --------
Mp_BAM2      --------
Sm_BAM2      --------
Ac_BAM2      --------
Eg_BAM2      --------
Rc_BAM2      --------
Fv_BAM2      --------
At_BAM2      --------
At_BAM1      --------
At_BAM3      --------
At_BAM6      --------
At_BAM5      --------
Ib_BAM5      --------
Tt_BAM   544 SVTITWQN


```
